# The enemy of my enemy is my friend: Nematode infection of pollinating and non-pollinating fig wasps has net benefits for the fig-fig wasp pollination mutualism

**DOI:** 10.1101/2020.05.08.084400

**Authors:** Justin Van Goor, Finn Piatscheck, Derek D. Houston, John D. Nason

## Abstract

Mutualistic associations between species pairs are ubiquitous in nature but are also components of broader organismal community networks. These community-level associations have shaped the evolution of individual mutualisms through interspecific interactions ranging from secondarily mutualistic to intensely antagonistic. Our understanding of this complex context remains limited because identifying species interacting with focal mutualists and assessing their associated fitness benefits and costs is difficult, especially over space and through time. Here, we focus on a community comprised of a fig and fig wasp mutualist, eight non-pollinating fig wasp (NPFW) commensals/antagonists, and a nematode previously believed to be associated only with the pollinator wasp mutualist. Through repeated sampling and field experiments, we identified that all NPFWs are targets for infection by this nematode. Further, this infection can impact NPFWs more severely than either mutualistic partner, suggesting a novel role of density-dependent facultative mutualism between fig and wasp mutualists and the nematode.

## Introduction

Mutualisms, or reciprocally beneficial interspecific interactions, are ubiquitous in nature and strongly influence ecological processes that, in turn, have shaped the trajectories of organismal evolution. Therefore, understanding the ecology and evolution of mutualistic associations is a crucial component to understanding ecosystem function. Much is understood regarding the co-evolutionary forces that are necessary for the formation and regulation of mutualisms (Axelrod and Hamilton 1981, Herre *et al* 1999, Lee 2015). Likewise, it is now widely understood that mutualisms can be context-dependent and can shift into commensalism and/or parasitism over time (Maynard Smith and Szathmary 1995, Bronstein 2001 and 2015, Song *et al* 2020). To date, the majority of theoretical (Ferriere *et al* 2002, Archetti 2019) and empirical (Heil *et al* 2009, Paterson *et al* 2010, Nelson *et al* 2018) studies have focused on the pairwise interaction between obligate mutualistic partners. Virtually all mutualistic species pairs, however, are members of more complex communities and networks of organismal interactions that may range from secondarily mutualistic, neutral, or strongly antagonistic in nature. This context is often lacking, but necessary for a clearer understanding of the ecological and evolutionary dynamics of mutualistic systems (Herrera *et al* 2002, Palmer *et al* 2010, Levine *et al* 2017, Chagnon *et al* 2020, Hall *et al* 2020).

Although mutualistic interactions are universal targets for antagonism through ecological roles such as predation, parasitism, and/or competition by community associates, the evolutionary consequences of such antagonisms are not widely understood. Generally, ecological theory predicts that interaction with community-level associates stabilizes or enhances mutualism fitness (Morris *et al* 2003, Jones *et al* 2009, Banerjee *et al* 2020, Chagnon *et al* 2020). However, theoretical works estimating negative (Ferriere *et al* 2002, Mougi and Kondoh 2014, Bachelot and Lee 2020) and neutral (Arizmendi *et al* 1996, Bronstein 2001) effects also exist, showcasing a presumable role for context-dependence in the diversity of community assemblages. The body of empirical research evaluating the role of community-interactions on mutualism fitness is limited, but growing (Bronstein 1991, Thompson and Fernandez 2006, Silknetter *et al* 2018, Song *et al* 2020). One impediment to investigating the effects of community-level antagonism on mutualism fitness is that lifetime fitness in many systems is difficult to quantify (West *et al* 1996, Bronstein 2001). This can be alleviated by focusing on model systems in which all intimately interacting species are known, ecological roles as mutualists and exploiters are well understood, and key components of lifetime fitness are easily estimated.

One such model system is the fig-fig wasp obligate pollination-nursery mutualism. Figs (Moraceae, *Ficus*) are represented by more than 750 species worldwide, with approximately 150 New World species (Berg 1989). Figs are ecologically important “keystone” components of tropical and subtropical ecosystems because their aseasonal fruit production serves as food resources for diverse assemblages of animals (Terborgh 1986, Shanahan *et al* 2001). *Ficus* species produce a nearly closed, urn-shaped inflorescence (a fig) attractive to wasps through volatile compounds (Chen *et al* 2009, Wang *et al* 2016). All Neotropical figs are monoecious (male and female flowers present in the same individual), and their inflorescences may contain tens to thousands of female and male flowers, varying among species (Herre 1989). Figs are entirely reliant on typically host-species-specific fig wasps (Hymenoptera: Agaonidae, many genera) for pollination services, and pollinator wasp larvae develop within a subset of the fig’s ovules (Janzen 1979). Pollinators have short adult life-spans (<60 hours; Kjellberg *et al* 1988, Dunn *et al* 2008a, Van Goor *et al* 2018), but excellent dispersal capabilities, exploiting wind currents to reach receptive, host-specific trees that are often located many kilometers from their natal trees (Nason *et al* 1998, Harrison and Rasplus 2006, Ahmed *et al* 2009).

In addition to obligate mutualistic relationships with pollinating wasps, individual species of *Ficus* are also subject to exploitation by a diversity of non-pollinating fig wasp (NPFW) genera (multiple Families) (Bouček 1993, Kerdelhué and Rasplus 1996, Farache *et al* 2018). Each fig species typically supports at least one, and often several, NPFW species (Compton and Hawkins 1992) and, like pollinating wasps, many are host-fig specific and appear to be attracted to receptive figs by the same volatile blends produced to attract pollinators (Proffit *et al* 2007). In contrast to the pollinator, which oviposits inside the fig, all Neotropical (and most Old World) NPFWs oviposit from the fig’s outer surface by inserting their ovipositors through the fig wall. Depending upon the species, NPFWs parasitize developing seeds, pollinators, or other non-pollinators, or induce galls within the fig wall in close proximity to developing pollinators (West *et al* 1996, Cruaud et al 2011, Segar *et al* 2018). Thus, many NPFW species have negative fitness impacts on the fig-pollinator mutualism (West and Herre 1994, Cardona *et al* 2013, Jansen-González *et al.* 2014, Borges 2015).

To date, the vast majority of research investigating antagonist effects on the fig-fig pollinator mutualism has focused on NPFWs (West *et al* 1996, Dunn *et al* 2008b, Elias *et al* 2012). Equally pervasive, but much less studied, are entomopathogenic nematodes that infect fig pollinators (Martin *et al* 1973). Nematodes of the genus *Parasitodiplogaster* (Diplogastridae) are pantropical associates of pollinating fig wasps (Poinar 1979, Poinar and Herre 1991). The life history of *Parasitodiplogaster* is tightly coupled with that of their pollinating wasp hosts, which they rely upon for energy, transport to a new fig, and subsequent reproductive success. For a description of the lifecycle of figs, their pollinator wasps, NPFWs, and *Parasitodiplogaster* see Figure 1. *Parasitodiplogaster* nematodes require transport to a new fig in each generation, and it is thus necessary that their impacts on female pollinator wasp survival are not so great as to prohibit successful dispersal to trees bearing receptive stage figs (Herre 1995, Van Goor *et al* 2018, Gupta and Borges 2019, Shi *et al* 2019). Despite this constraint, the virulence of nematode infection varies across species as a function of host-wasp species population density (Herre 1993) and can range from avirulent or commensal (Herre 1995, Ramirez-Benavides and Salazar-Figueroa 2015, Van Goor *et al* 2018, Shi *et al* 2019) to virulent (Herre 1993, 1995), reducing host offspring production by up to 15%.

**Figure 1.**
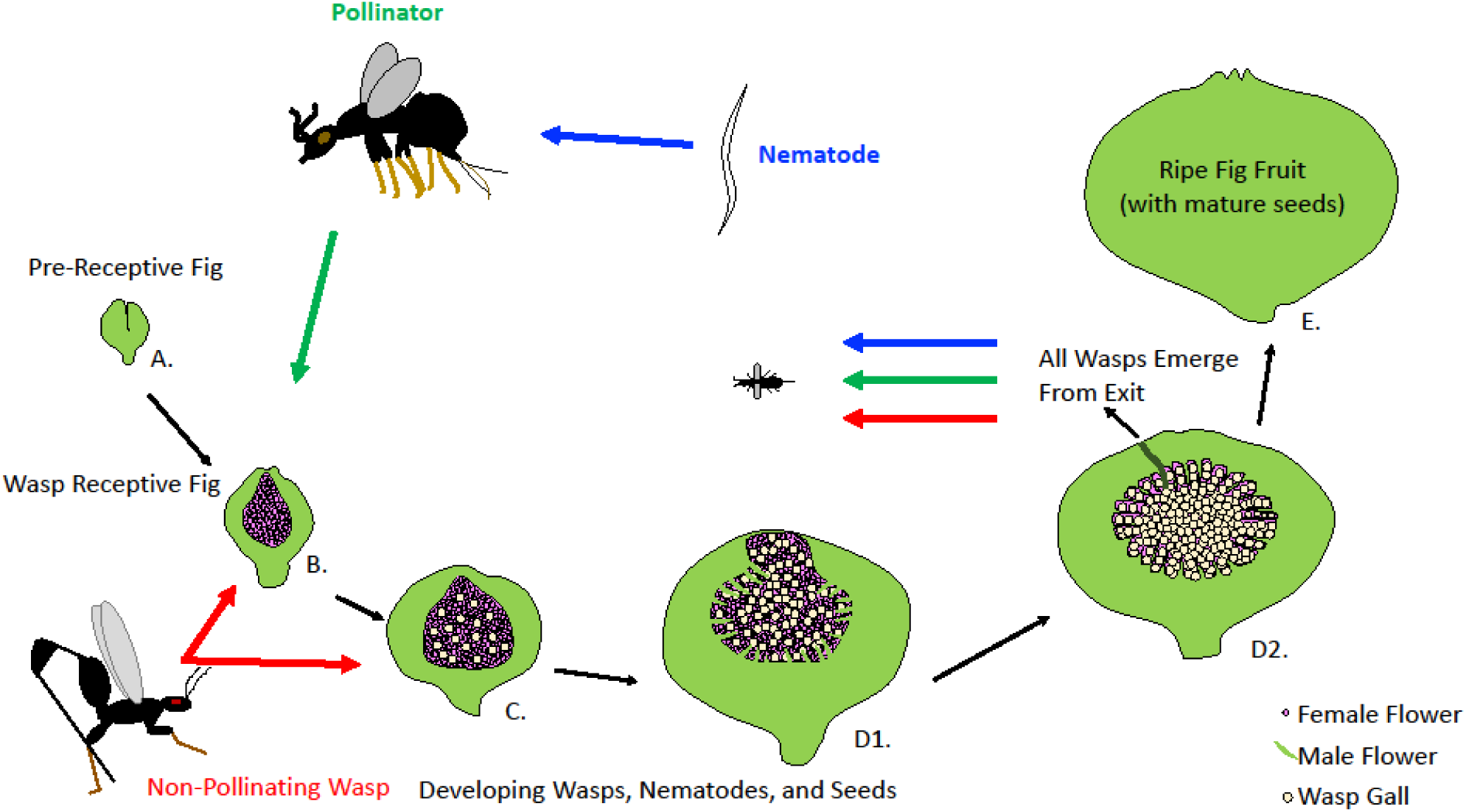
Parallel lifecycles of monoecious figs, pollinating fig wasps, non-pollinating fig wasps (NPFW) and *Parasitodiplogaster* nematodes. Pre-receptive phase figs (A) grow on *Ficus* branches before the development of internal female flowers. The fig is then receptive to pollinating wasps (B), which enter the fig through a terminal pore (ostiole) and then pollinate a subset of flowers and oviposit eggs into another subset before dying. If the pollinator was infected by nematodes they will molt from infective juveniles to consumptive adults that feed on wasp tissue before molting again into reproductive adults. They will then aggregate outside of the pollinator and form mating clusters before female nematodes disperse throughout the fig to lay eggs and then die. During (B) and early (C) phases NPFW species oviposit eggs from the exterior of the fig into developing wasp galls, unpollinated florets, or developing seeds. During the interfloral phase (C) pollinator and NPFW offspring larvae develop in galls, nematode eggs develop on these galls, and seed development takes place. In early male-phase (D1), male flowers develop within the fig, adult male pollinators and NPFWs emerge to inseminate females and release them from their galls, and infective stage juvenile nematodes position themselves on wasp galls. In later male-phase (D2) female pollinators collect pollen from flowers and juvenile nematodes perform nictation behavior to contact and infect hosts. Using a hole bored by pollinator males, female pollinators, NPFWs, and nematodes all exit the fig to start the cycle anew. As fig seeds reach maturity (E) the fig becomes pumped with sugar to promote herbivory and seed dispersal. Total developmental time for the *F. petiolaris* community (A-E phases) is between six and ten weeks (Van Goor and Piatscheck *personal observation*), depending on season. Image modified and re-drawn from Galil and Eisikowitch 1968.

While infection by *Parasitodiplogaster* nematodes can negatively influence the fitness of pollinating fig wasps (definitive hosts), the incidence and fitness effects of nematode infection on co-occurring NPFWs has not been described. Although pollinator and NPFWs share the same developmental space within the fig and are both exposed to infective juvenile nematodes while emerging from a mature fig, infection of NPFWs should be maladaptive for the nematodes. Pollinator wasp hosts enter figs to lay their eggs, granting infective nematodes access to the next generation of emerging hosts. Conversely, all NPFWs oviposit from the exterior of the fig (Bouček 1993, Elias *et al* 2008), precluding associated nematodes access to the interior of the fig and new hosts. Therefore, nematode infection of NPFWs has been described as a maladaptive behavior that should be strongly selected against (Giblin-Davis *et al* 1995, Vovlas and Larizza 1996, Krishnan *et al* 2010). Surprisingly, however, *Parasitodiplogaster* has been reported to infect multiple NPFW species (Van Goor *et al* 2018), and has been observed in a number Panamanian fig-host communities as well (Van Goor, *personal observation*). If nematodes negatively impact the fitness of NPFWs that interact antagonistically with figs and their pollinators, they could have previously unappreciated benefits for fitness and persistence in the fig-pollinator mutualism.

Here, we investigate how *Parasitodiplogaster* nematode infection may limit NPFW fitness and, in turn, potentially benefit the mutualistic partnership between figs and wasps (Figure 2). Utilizing the Sonoran Desert rock fig, *Ficus petiolaris*, as a study system, we document variation in nematode-NPFW interactions across geographic locations and quantify the fitness impacts of nematode infection on eight NPFW species. First, we quantify the level of antagonism between NPFW species and the fig-pollinator mutualists. We then determine the incidence and number of nematodes infecting NPFWs. Finally, we quantify the impacts of infection on successful NPFW dispersal ability to estimate disruptions or limitations in life history.

**Figure 2.**
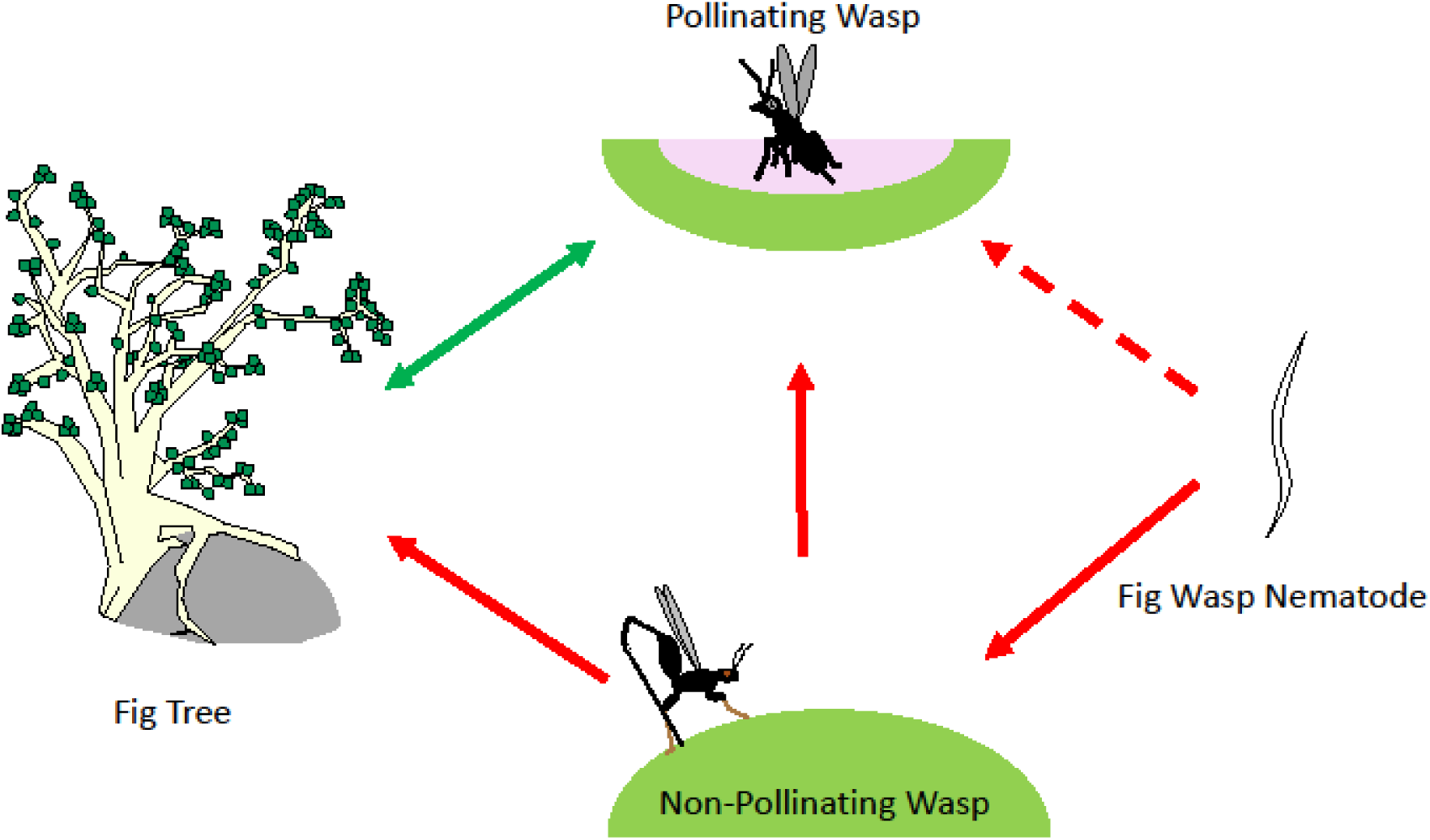
Hypothesized interaction schematic between figs, pollinating fig wasps, non-pollinating fig wasps (NPFW), and *Parasitodiplogaster* nematodes. The solid green arrow indicates the well-known mutualism between figs and their pollinating fig wasps. NPFWs have demonstrable antagonistic and fitness-reducing effects (solid red arrows) against both figs and pollinating fig wasps, depending on species. Fig wasp nematodes can range from virulent to relatively benign against pollinator wasp hosts. *Parasitodiplogaster* nematodes associated with *Pegoscapus* pollinators of *F. petiolaris* have been shown to be largely benign in their virulence against their hosts (dotted red arrow). The interaction between fig nematodes and NPFWs is not understood but is hypothesized to be antagonistic (solid red arrow) due to the average length of time infective nematodes would likely be associated with these wasps. Further, it is hypothesized that nematode-induced reduction of NPFW may directly benefit pollinating wasps and their host fig trees.

## Material and Methods

### *The* Ficus petiolaris *system of Northwestern Mexico*

*Ficus petiolaris* is a monoecious rock-strangling fig that is widespread in the desert and seasonally-xeric habitats of Baja California and mainland Mexico. The nine census sites investigated in this study are located in the states of Baja California and Baja California Sur (Supplemental Table and Figure 1), where *F. petiolaris* is the only native fig species. *Ficus petiolaris* is obligately pollinated by an unclassified *Pegoscapus* wasp (Agaonidae), which appears to be a single species based on phylogenetic analyses (Su *et al* 2008, Satler *et al* 2019). *Pegoscapus* wasps die inside figs after pollination and oviposition, and the number of foundress wasps contributing offspring to each fig can counted. This *Pegoscapus* species is subject to parasitism by a single species of *Parasitodiplogaster* nematode, whose 28S rDNA sequences form a single, well-supported clade that clusters with other publicly available Neotropical *Parasitodiplogaster* sequences (Van Goor *et al* 2018). No other fig-associated nematode genera (*Schistonchus, Pristionchus, Ficophagus, Caenorhabditis*; Vovlas and Larizza 1996, Susoy *et al* 2016, Davies *et al* 2017, Woodruff and Phillips 2018) have been observed in *F. petiolaris* figs.

In addition to a pollinator wasp and associated *Parasitodiplogaster, F. petiolaris* in Baja California is host to eight chalcidoid NPFW species. This community is comprised of three *Idarnes* species (Sycophaginae, Munro *et al* 2011, Cruaud *et al* 2011, Satler *et al* 2020), one from species group *flavicollis* (ovule-gallers) and two from species group *carme* (kleptoparasites or parasitoids sensu Elias *et al* 2012, Farache *et al* 2018). These three species are referred herein as *Idarnes flavicollis* and *Idarnes carme* species 1 and 2, respectively. Additionally, there are two species of *Heterandrium* (Pteromalidae), both of which gall *F. petiolaris* ovules (Duthie and Nason 2016). These five species are the most commonly observed NPFWs associated with *F. petiolaris*, and as ovule-gallers, kleptoparasites, and/or seed parasites either compete directly with *Pegoscapus* pollinators for reproductive resources or utilize them for their own development (Duthie and Nason 2016). *Ficus petiolaris* is also host to one species of *Ficicola* (Pteromalidae) that generates large galls protruding from the receptacle into the interior of the fig and which may spatially impact developing seeds or larvae (Conchou *et al* 2014). Finally, one species of *Physothorax* (Torymidae) and one species of *Sycophila* (Eurytomidae) are parasitoids that develop within other fig wasp larvae (West *et al* 1996, van Noort *et al* 2013, Farache *et al* 2018).

### *Which NPFWs Antagonize the* F. petiolaris *Mutualism?*

Mature *F. petiolaris* trees were geo-referenced at nine sites along a latitudinal gradient spanning 741 km of the Baja California peninsula. Mature, wasp-releasing figs were sampled from each site, measured, pollinating foundress wasps counted, pollinating and NPFWs offspring produced per fig collected, and the presence/absence of juvenile nematodes assessed. These study sites were visited at four time points (November-December 2012, May-July 2013, November-December 2013, and May-July 2014) to ensure adequate sample sizes of wasp producing figs in both wet (October-December) and dry (May-July) seasons. The wasp offspring were preserved in 95% ethanol and figs were air-dried and placed into coin envelopes. Pollinating and NPFWs per fig were tallied by species and sex. A subset of the dried figs were cut into quarters and suspended in 90% ethanol to determine seed production.

We analyzed the effects of NPFWs and nematodes on two primary fitness components of the mutualism: pollinator offspring and seed production per fig. We used a Generalized Linear Mixed Model (GLMM) with Poisson errors and a log-link function to study *pollinator offspring per fig* as a response variable against the predictor variables *tree* nested within *site, season* (wet or dry), *pollinator foundress count, fig volume* (*mm*^*3*^), *nematode infestation* (presence or absence within the fig), and the *number of NPFW offspring* produced by each of the eight NPFW species. These GLMM analyses were conducted using the `glmer` function in the `lme4` package (Bates *et al* 2015) for R (R Core Team 2018).

We used other GLMMs to reciprocally study *seed production per fig* as well as the offspring production of each NPFW as response variables against the same predictor variables as in the preceding model. The directionality and significance of association observed between species offspring production in these models can allow for inference of ecology and potential antagonism against figs and pollinators, but should be viewed in the appropriate ecological context (Raja *et al* 2015). Significant positive associations can predict kleptoparasitism or parasitoidy, while significant negative associations could suggest competition for resources. In all models, the predictor variables *tree* nested within *site* and *season* were treated as random effects because of the over-dispersion of pollinator offspring and seeds observed between trees within sites over time.

### Frequency of Interaction between Nematodes and NPFWs

*Parasitodiplogaster* nematodes associated with *F. petiolaris* infect an ecologically relevant proportion of pollinating wasps in each generation (39% across sites; Van Goor *et al* 2018). Surprisingly, *Parasitodiplogaster* nematodes have been found to commonly infect NPFWs in *F. petiolaris* (Van Goor *et al* 2018) and in multiple Panamanian fig communities (Van Goor, *personal observation*). Focusing here on nematode infection of the *F. petiolaris* NPFWs, we sampled wasps of each NPFW species emerging from mature nematode infested figs across each study site and throughout time. The thoracic and abdominal cavities of these wasps were dissected using 0.25mm diameter tungsten needles (Fine Science Tools®) to determine the presence and number of infective juvenile nematodes.

### Nematode Infection Effects on NPFW Life History

An effective inference of life history reduction due to nematode infection can be developed through an evaluation of dispersal ability limitation. Only high levels of *Parasitodiplogaster* infection (> 10 individuals per host) have been associated with reduced dispersal in the pollinating fig wasps of *F. petiolaris*, leading to the estimated loss of 2.8% of pollinating wasp hosts per generation (Van Goor *et al* 2018). Similar reductions in dispersal ability could occur in NPFWs infected with nematodes, but this has yet to be investigated. To estimate the effect of nematode infection on NPFW dispersal ability, the number of nematodes infecting NPFWs emerging from mature, nematode infested figs was compared to the number of nematodes infecting successfully dispersed NPFWs arriving at receptive figs. Successfully dispersed wasps were collected from receptive figs from two sampling sites (70 and 96) by J. Nason and J. Stireman in 2004-2005 and from Site 96 by J. Van Goor in August 2016. These wasps were preserved in 95% ethanol for dissection. If nematode infection decreases NPFW dispersal, then we predict that successfully dispersed wasps will contain fewer nematodes than wasps just emerging from nematode infested figs. The comparison of infection rates between emerging and successfully dispersed wasps of each NPFW species was conducted by exact test using the `poisson.test` function in R.

## Results

### *Which NPFWs Antagonize the* F. petiolaris *Mutualism?*

The four field collections conducted from 2012 to 2014 yielded a total of 2187 mature, wasp-producing figs. We obtained an average of 260.1 figs (range 166-373) per site, with each fig producing an average of 84.7 (range of 2 to 503) fig wasps. Of these wasps, 40.1% (36.5 per fig) were pollinators and 59.9% (48.2 per fig) were NPFWs. All eight NPFW species were observed at all sites except for *Physothorax* and *Sycophila*, which were absent at one and two sites, respectively. The three *Idarnes* species were the most abundant NPFWs, collectively accounting for an average of 49.6% of all wasps observed across the *F. petiolaris* sites, with *Idarnes flavicollis* as the most common (24.2% of all wasps, Supplemental Table 2). Illustrative of the high abundance NPFWs in the *F. petiolaris* community, we observed only 15 (0.69%) figs in which NPFWs were absent. Interestingly, 160 (7%) of the figs surveyed contained zero pollinating foundresses and no pollinating offspring, yet still produced NPFW offspring. Nematode infestation was not observed in any of these zero-foundress figs.

Potential antagonistic effects of NPFWs on pollinator offspring and fig seed production were evaluated using GLMM analyses. In the pollinator model (GLMM, *n* = 2187, *df* = 11, Log Likelihood = −29544.2), we found *pollinator offspring per fig* to be significantly associated with *fig volume* (*mm*^*3*^), *pollinator foundress count*, and *nematode infestation* (all *p*-values < 0.001). Figs infested with nematodes produced significantly fewer pollinator offspring, but this limitation only accounted for a 1% reduction (mean of 36.80 pollinator offspring in infested figs and 37.17 in uninfested). Interestingly, pollinator offspring production had a range of significant (both positive and negative) and non-significant associations with NPFWs (Table 1).

**Table 1.** Matrix indicating the result of GLMM analyses showcasing the reproductive output of members of the *F. petiolaris* community of northwestern Mexico and subsequent associations between member species. The analysis of *F. petiolaris* seed production represents a subset of the total dataset. + indicates a significant positive association (*p* = < 0.05), - indicates a significant negative association (*p* = < 0.05), and ns indicates non-significance (*p* = > 0.05).

|  | Fig<br>Seeds | *Pegoscapus* | *Idarnes*<br>*flavicollis* | *Idarnes*<br>*carne 1* | *Idarnes*<br>*carne 2* | *Heterandrium*<br>*1* | *Heterandrium*<br>*2* | *Ficicola* | *Phsyothorax* | *Sycophila* |
| --- | --- | --- | --- | --- | --- | --- | --- | --- | --- | --- |
| **Fig Seeds** | / |  |  |  |  |  |  |  |  |  |
| *Pegoscapus* | + | / |  |  |  |  |  |  |  |  |
| *Idarnes*<br>*flavicollis* | - | - | / |  |  |  |  |  |  |  |
| *Idarnes*<br>*carne 1* | **ns** | + | - | / |  |  |  |  |  |  |
| *Idarnes*<br>*carne 2* | + | + | - | + | / |  |  |  |  |  |
| *Heterandrium*<br>*1* | **ns** | - | + | - | + | / |  |  |  |  |
| *Heterandrium*<br>*2* | + | + | + | + | + | + | / |  |  |  |
| *Ficicola* | + | **ns** | + | **ns** | **ns** | **ns** | **ns** | / |  |  |
| *Phsyothorax* | - | - | - | **ns** | + | **ns** | **ns** | + | / |  |
| *Sycophila* | **ns** | - | **ns** | **ns** | **ns** | **ns** | **ns** | + | **ns** | / |

Of the mature, wasp producing figs collected, a total of 120 (60 each from two sites) were examined to investigate potential antagonistic effects of NPFWs with fig seed production. The fig seed model (GLMM, *n* = 120, *df* = 12, Log Likelihood = −827.20) revealed a highly significant relationship between *seed production per fig* and the predictor variable *fig volume* (*mm*^*3*^) (*p*-value < 0.001). Although seed production was not significantly associated with *pollinator foundress count* (*p* = 0.746), it had a significantly positive association with *pollinator offspring production* (*p* = 0.048). Surprisingly, seed production was found to be significantly *higher* (by 10.3%) in figs with *nematode infestation* (*p* < 0.001). As in the pollinator GLMM, significant associations between NPFWs and fig seed production ranged from positive to negative, while others were not significant (Table 1). *Idarnes flavicollis* and *Physothorax* were significantly negatively associated with *both* pollinator offspring and fig seed production. Conversely, *Idarnes carme* 2 and *Heterandrium* 2 was significantly positively associated with both of these reproductive measurements.

### Frequency of Interaction between Nematodes and NPFWs

*Parasitodiplogaster* nematode infestation was observed in 36% (780 of 2187) of all mature figs sampled, varying between 12 and 80% depending on individual study site and collection trip. From these figs, a total of 2791 emerging pollinators and NPFWs (range of 4 to 1182 individuals depending on species) were dissected to determine the presence and number of infective juvenile nematodes. With the exception of *Sycophila* (presumably due to low sample size), all NPFW species were parasitized by *Parasitodiplogaster*, with the incidence of infection in individual wasps varying substantially among host species, ranging from 6.7 to 39.6% (Table 2). Interestingly, the number of nematodes per infective event also varied substantially among NPFW species. *Idarnes flavicollis* and *Heterandrium* 1, in particular, experienced high incidences of nematode infection (39.6% and 27.8% respectively) and also high average nematode loads (2.52 and 2.32 nematodes per host) that were significantly greater than in other NPFW species, though not as high as in *Pegoscapus* pollinators (Supplemental Table 3).

**Table 2.** Parasitism of *F. petiolaris* associated pollinator (*Pegoscapus*) and NPFW (all other genera) wasp species by *Parasitodiplogaster* nematodes. Results are from wasps emerging from mature, nematode-infested figs and are pooled across all collections.

| Wasp Species | Wasps Dissected | Percent Infected | Average Nematodes Per Host | Median Nematodes Per Host | Maximum Nematodes Per Host |
| --- | --- | --- | --- | --- | --- |
| *Pegoscapus* | 1182 | 61.5% | 4.03 | 3 | 50 |
| *Idarnes flavicollis* | 573 | 39.6% | 2.52 | 2 | 21 |
| *Idarnes carme* sp. 1 | 460 | 8.5% | 1.21 | 1 | 4 |
| *Idarnes carme* sp. 2 | 135 | 13.3% | 1.56 | 1 | 5 |
| *Heterandrium* sp. 1 | 110 | 27.8% | 2.32 | 1 | 15 |
| *Heterandrium* sp. 2 | 122 | 18.8% | 1.65 | 1 | 4 |
| *Ficicola* | 104 | 6.7% | 1.57 | 1 | 4 |
| *Physothorax* | 59 | 8.5% | 1.60 | 1 | 4 |
| *Sycophila* | 4 | 0% | N/A | N/A | N/A |

### Nematode Infection Effects on NPFW Life History

A total of 281 NPFWs of various species were collected as they were arriving at receptive fig trees in 2004-2005 (146 individuals) and August 2016 (135 individuals). Although we were able to sample six of the eight NPFW species associated with *F. petiolaris*, many of these were sampled at very low densities (Supplemental Table 4). However, *Idarnes flavicollis* and *Idarnes carme* 1 and 2 were all collected more frequently, with sample sizes of 111, 86, and 71 individuals, respectively. Of these three species, very few were observed arriving at receptive fig trees infected with juvenile nematodes (3.6%, 1.2%, and 2.8%, respectively). Even within these infected wasps, it was uncommon for the number of juvenile nematodes to exceed one individual. This effect was only observed *once* with one *Idarnes flavicollis* wasp carrying two juvenile nematodes. Indeed, in each of these three cases, there were significantly fewer individual wasps arriving with nematodes and fewer infective nematodes per individual compared to those wasps leaving nematode infested figs (poisson.test, *p*-values = < 0.001, 0.005, and < 0.001 respectively). Thus, of these three NPFW species we found that both the rate of infection and the number of nematode individuals observed per infective event are lower in wasps that successfully dispersed to receptive figs compared to those emerging from infested figs (Table 3).

**Table 3.** *Idarnes* NPFWs infected with *Parasitodiplogaster* nematodes appear unlikely to successfully disperse to receptive figs. Presented here is the dispersal information for three abundant NPFW species that were collected arriving at receptive fig trees. First, we report the percent of individuals emerging from mature figs infected with at least one juvenile nematode (from Table 2). Next, given that we found 36% of all figs to be infested with nematodes from our general dataset, we estimate the total percentage of individuals infected with nematodes at each generation. We compared this to the total number of successfully dispersed individuals that were found infected by at least one juvenile nematode at receptive fig trees. To estimate the rate of failure to disperse for these wasps we divided the percent of individuals arriving with nematodes from those emerging with nematodes. Finally, to estimate the total number of NPFW individuals removed from the population due to nematode infection in each generation, we multiplied the total percent of individuals infected in the population by the appropriate failure to disperse rate.

| Non-pollinator wasp species | Percent infected individuals emerging from infested fig | Percent of total non-pollinator population infected | Percent infected individuals arriving at receptive fig | Failure rate of infected fig wasp to arrive at receptive fig | Total percent of non-pollinator individuals excluded due to nematode infection |
| --- | --- | --- | --- | --- | --- |
| *Idarnes flavicollis* | 39.6% | 14.3% | 3.6% | 0.91 | 13% |
| *Idarnes carme* species 1 | 8.5% | 3.1% | 1.2% | 0.86 | 2.7% |
| *Idarnes carme* species 2 | 13.3% | 4.8% | 2.8% | 0.79 | 3.8% |

## Discussion

### *Which NPFWs Antagonize the* F. petiolaris *Mutualism?*

Like the vast majority of fig systems, the NPFW (and nematode) community associated with *F. petiolaris* is speciose (Bouček 1993, Marussich and Machado 2007, Castro *et al* 2015). Interestingly, the total assemblage of *F. petiolaris* associated NPFWs typically outnumber pollinating wasp mutualists, which is partially explained by the density of host trees in which this fig community occurs (Duthie and Nason 2016). Surprisingly, we observed that 7% of all figs surveyed produced NPFW offspring without the presence of a pollinating foundress. In fact, all NPFW genera associated with *F. petiolaris* were produced in these figs. The fact that *Parasitodiplogaster* nematodes were not observed in any of these zero-foundress figs reinforces that NPFW species are not reliable vectors for nematode transmission to receptive figs, as previously suggested (Giblin-Davis *et al* 1995, Vovlas and Larizza 1996, and Jauharlina *et al* 2012).

Within the NPFW community of *F. petiolaris*, three wasps of the genus *Idarnes* were notably common, consisting of nearly 50% of all wasps collected. This abundance is consistent with observations in other Neotropical fig communities (West *et al* 1996, Elias *et al* 2012, Farache *et al* 2018), making them extremely relevant antagonists for community-level evaluation. Of the *F. petiolaris*-associated *Idarnes, Idarnes flavicollis* was particularly abundant, accounting for 24% of all wasps sampled (Supplementary Table 2). Indeed, in the GLMM analyses that were performed, we found strong negative associations between *Idarnes flavicollis* offspring production and pollinator offspring production *as well as* fig seed production (Table 1), suggesting a critical antagonistic role of this particular species against this mutualism. Similar significant negative associations were observed in *Physothorax* (both seeds and pollinators), *Heterandrium* 1, and *Sycophila* (just pollinators). These NPFWs are predicted to directly compete for the same reproductive resources utilized by the mutualist-partners through spatial interruption or exclusion of developing fig seeds or pollinator larvae, as discussed in Conchou *et al* (2014). Other species indicated significant positive associations against pollinator offspring production and/or seed production even when accounting for other important biological and geographic variables, suggesting ecologies such as kleptoparasites or parasitoids (Raja *et al* 2015). Interestingly, many NPFW did not have significant associations with each other (presumably due to limited sample sizes) except for the positive association of *Ficicola* and *Physothorax*, which have a known host-parasitoid relationship (Bouček 1993).

Importantly, these GLMMs also suggest that nematode infection, while significantly negatively associated with pollinator wasp offspring production, is a relatively benign effect that only limits an estimated 1% of the pollinator population every generation even though this infection is relatively common (consistent with Van Goor *et al* 2018, Gupta and Borges 2019, and Shi *et al* 2019). In *F. petiolaris*, we have previously observed that nematode infection does not appear to limit pollinator longevity, dispersal ability, or offspring production unless many nematode individuals attempt to infect the same host (Van Goor *et al* 2018). Surprisingly, we have also found a significant positive association between nematode infestation and seed production (by 10%). This effect has not been previously observed in figs with *Parasitodiplogaster* nematodes and the mechanism through which this increased seed production takes place is not currently understood. Thus, these GLMM analyses incorporating important geo-spatial and biological information provides some speculation as to the notable NPFW antagonists against the *F. petiolaris* mutualism (*Idarnes* and *Heterandrium*) but also strongly suggests a benign (or even slightly beneficial) role for *Parasitodiplogaster* nematodes.

### Frequency of Interaction between Nematodes and NPFWs

Intriguingly, nearly all NPFW species associated with *F. petiolaris* were found to be subject to infection by *Parasitodiplogaster* nematodes (Table 2). This effect is unexpected because this infection represents a reproductive dead-end for the nematodes. Nevertheless, nematode infection of NPFWs occurs frequently, ranging from 6-40% of all individuals leaving nematode infested figs, depending on the wasp species. This effect should be strongly selected against (Krishnan *et al* 2010) but could be explained by the crowded and fast-moving milieu in which juvenile nematodes actively attempt to contact a pollinating wasp host (Figure 1). Pollinating and NPFW hosts emerge into the same cramped environment and leave the host fig within minutes to hours. Perhaps nematodes have not evolved to become choosy in this situation and instead infect the first wasp they contact through nictation. Additionally, most figs in *F. petiolaris* are visited by a single pollinating foundress (Duthie and Nason 2016, Van Goor *et al* 2018), which will likely only introduce a single lineage of nematodes, if infected. Infection of NPFWs instead of definitive pollinating wasp hosts could be explained through kinship theory (Hamilton 1964), but the genetic data needed to support this hypothesis is not currently available.

Pollinating fig wasps are the “appropriate” hosts for *Parasitodiplogaster* nematodes because these wasps enter into receptive figs and secure reproductive space for nematodes. Indeed, we observed here that pollinators are infected more frequently (Table 2) and with significantly higher nematode loads (Supplemental Table 3) than NPFW hosts. However, certain NPFW species are infected more frequently and tend to have higher nematode loads than other NPFWs. Interestingly, this appears to correspond with the NPFW species that are most abundant and have the most fitness-limiting effects for the mutualism (*Idarnes flavicollis, Physothorax*, and *Heterandrium* 1). Nematodes that infect pollinators should delay their potentially fitness-limiting behavior until their pollinator host wasp has successfully arrived at a receptive fig so that they can better ensure their own reproductive opportunities. Pollinating fig wasps are relatively short lived (typically < 60 hours, Kjellberg *et al* 1988, Van Goor *et al* 2018), meaning that nematode-molting cues (Figure 1) have likely evolved in response. However, NPFWs have much longer life histories (mean 150-350 hours depending on species, unpublished data) in which they search for receptive figs or oviposit into multiple figs (Ghara *et al* 2014). NPFWs infected with nematodes thus spend significantly more time in this association, which may have severely detrimental effects on NPFW longevity, dispersal ability, and therefore overall fitness.

### Nematode Infection Effects on NPFW Life History

Efforts were made to conduct NPFW longevity trials (similar to those presented in Van Goor *et al* 2018) but failed to show significant effects of nematode infection on NPFW longevity (Supplemental Tables 5 and 6). This was likely due to low NPFW sample sizes and to the closed conditions in which the trials took place (small plastic vials). Notably, all participant wasps remained relatively stationary within these vials until they died. These conditions do not appropriately represent the natural stresses presented to NPFW wasps that typically must disperse, oviposit their eggs, and avoid predation throughout their lifespans, and should therefore be treated with interpretive caution.

NPFWs that fail to arrive at receptive figs due to nematode infection are incapable of reproducing, eliminating their individual fitness. To more naturally estimate the effects of nematode infection on dispersal ability we compared the frequency of infection and the number of nematode individuals involved per infective event for NPFWs that were emerging from nematode infested figs to those that had successfully dispersed to receptive figs. Pollinating wasp mutualists of *F. petiolaris* tolerate moderate levels of nematode infection (< 10 individuals) without any correlated reductions in dispersal ability or offspring production (Van Goor *et al* 2018). Alternatively, we infrequently observed *F. petiolaris*-associated NPFWs successfully arriving at receptive figs with *any* nematode infection. When infection was observed (in only 7 of 275 wasps) it was typically with only a single nematode, and only one instance in which two nematodes were observed (Supplemental Table 4). This strongly suggests that NPFW antagonists of *F. petiolaris* do not have the same tolerance to nematode infection that pollinating mutualists experience, and that nematode infection severely limits NPFW dispersal ability and reproductive capabilities in natural environments.

We previously estimated that pollinators lose 2.8% of the general population each generation due to nematode overexploitation (Van Goor *et al* 2018). Here, we have identified that each NPFW associated with *F. petiolaris* is target for the same infection and are likely more sensitive to infection by even a single nematode. Looking into the abundantly available arriving NPFWs from the genus *Idarnes*, we can similarly estimate net losses due to infection that are similar or substantially higher than those suffered by pollinators each generation (Table 3). *Idarnes flavicollis*, which has been identified as the *most abundant* and one of the *most antagonistic* NPFWs associated with *F. petiolaris* loses an estimated 13% of the general population due to nematode infection every generation. It is likely (with increased sampling) that all other associated NPFW suffer fitness limitations due to infection, suggesting that nematode infection may remove a sizeable proportion of the total wasp-antagonist community in each generation. This may be explained as an indirect effect (Gillespie and Adler 2013, Guimarães *et al* 2017) or may represent a novel density-dependent facultative mutualism between *Parasitodiplogaster* nematodes, *Pegoscapus* pollinating wasps, and *F. petiolaris*. When more NPFWs exploit the mutualism they are more likely to encounter nematodes whose infection they are sensitive to, making them unlikely to arrive at a new fig and reproduce. Ultimately, this may present a mechanism through which antagonist communities are modulated over shared community resources.

### Conclusion

*Parasitodiplogaster* infection of NPFW associated with *F. petiolaris* is widespread and likely has substantial ecological and evolutionary consequences for fig community dynamics. Similar NPFW infection has been observed in five other Panamanian fig communities, suggesting that this phenomenon is much more widespread than previously recognized (Van Goor, *personal observation*). In particular, NPFWs associated with *F. popenoei* may be infected more frequently than pollinators (Supplemental Table 7) and may experience more detrimental fitness limitations (Supplemental Tables 8 and 9). In addition, *Parasitodiplogaster* nematodes may be able to clear figs of harmful and widespread *Fusarium*-like fungal infections that are capable of eliminating entire crops from fig trees (Michailides *et al* 1996). The mechanism underlying this secondarily mutualistic behavior of *Parasitodiplogaster* for fig systems is currently unknown, but will be the target of future research. These findings together suggest a more complex set of interactions underlying community-level dynamics that may be present beyond fig-systems. “Accidental infections” or similar facultative mutualisms may potentially act as a hidden driver for community dynamics, especially in lesser-studied, invertebrate-rich assemblages.

## Acknowledgements

The authors give thanks to N. Davis, A. Gómez, A. Gutiérrez, and A. Oldenbeuving for their assistance with field collections, and the dozens of undergraduate research assistants that helped identify and quantify wasp offspring from more than 2000 figs. We specifically thank T. Mores, S. Pahlke, N. Rohlfes, S. Skinner, and L. Thiesse for their assistance with wasp dissections. We also wish to thank EA. Herre for his thoughtful insight with the development of this manuscript. This research was supported by funding from the National Science Foundation (awards DEB-0543102 to J. Nason and R. Dyer, DEB-1146312 to J. Nason, and DEB-1556853 to J. Nason, T. Heath, and EA. Herre), the University of Iowa Center for Global and Regional Environmental Research (to J. Van Goor), and the Iowa State University Department of Ecology Evolution and Organismal Biology Gilman Scholarship (to J. Van Goor).

## Supplemental Tables

**Supplemental Table 1.**
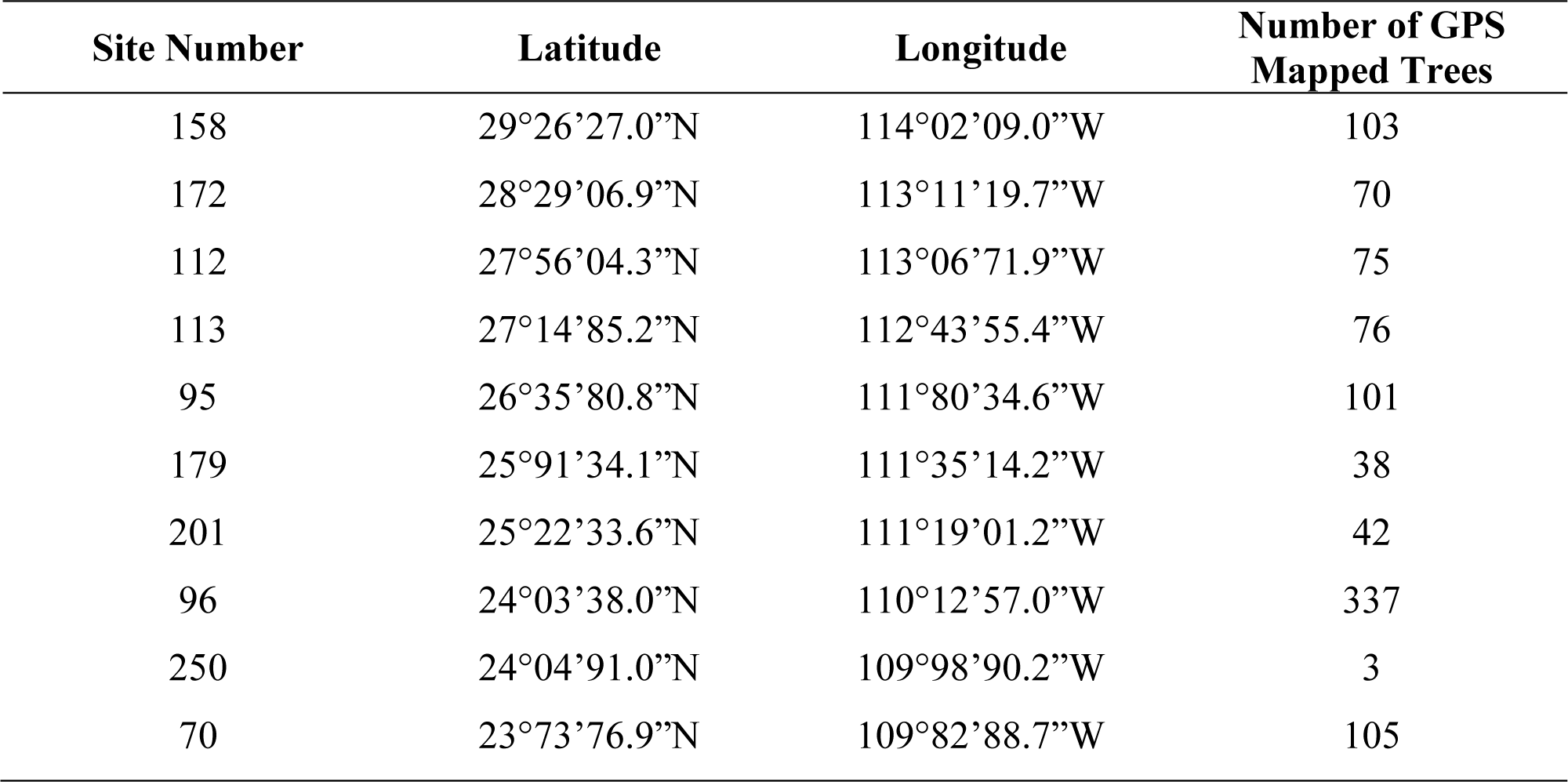
*Ficus petiolaris* study site identification number, latitude and longitude coordinates, and number of GPS-mapped trees.

**Supplemental Table 2.** The average number and percentage of pollinating and NPFW individuals reared per fig at each *F. petiolaris* study site in Baja California, Mexico. Sites are arranged from northernmost (158) to southernmost (70) in the table.

|  | *Pegoscapus* | *Idarnes*<br>*flavicollis* | *Idarnes*<br>*carne*<br>*sp.1* | *Idarnes*<br>*carne*<br>*sp. 2* | *Heterandrium*<br>*sp. 1* | *Heterandrium*<br>*sp. 2* | *Ficicola* | *Physothorax* | *Sycophila* |
| --- | --- | --- | --- | --- | --- | --- | --- | --- | --- |
| **158** | 13.2<br>(24.7%) | 14.5<br>(35.5%) | 4.2<br>(10.3%) | 8.5<br>(20.8%) | 1.4<br>(3.4%) | 0.8<br>(1.9%) | 0.2<br>(0.5%) | 0.5<br>(1.3%) | 0.6<br>(1.5%) |
| **172** | 39.7<br>(37.8%) | 16.6<br>(18.9%) | 9.8<br>(11.2%) | 18.4<br>(21.0%) | 4.1<br>(4.7%) | 3.1<br>(3.5%) | 1.0<br>(1.0%) | 1.7<br>(1.9%) | 0.02<br>(0.02%) |
| **112** | 24.8<br>(30.1%) | 17.3<br>(26.2%) | 5.3<br>(8.1%) | 13.1<br>(19.8%) | 3.6<br>(5.4%) | 5.2<br>(7.9%) | 1.0<br>(1.5%) | 0.5<br>(0.7%) | 0.1<br>(0.1%) |
| **113** | 27.0<br>(32.6%) | 8.8<br>(13.1%) | 6.8<br>(10.1%) | 21.4<br>(31.9%) | 3.1<br>(4.6%) | 2.0<br>(3.0%) | 1.0<br>(1.5%) | 2.0<br>(3.0%) | 0.1<br>(0.1%) |
| **95** | 55.5<br>(52.7%) | 18.4<br>(20.0%) | 7.3<br>(7.9%) | 8.5<br>(9.2%) | 4.7<br>(5.1%) | 1.6<br>(1.8%) | 0.9<br>(1.0%) | 2.0<br>(2.2%) | 0.01<br>(0.01%) |
| **179** | 16.4<br>(21.5%) | 18.5<br>(32.5%) | 9.5<br>(16.8%) | 11.9<br>(21.1%) | 0.8<br>(1.4%) | 2.2<br>(4%) | 1.3<br>(2.3%) | 0.1<br>(0.2%) | N/A |
| **201** | 26.9<br>(29.5%) | 33.8<br>(46.4%) | 3.2<br>(4.4%) | 7.4<br>(10.2%) | 3.8<br>(5.2%) | 0.6<br>(0.9%) | 0.8<br>(1.1%) | 1.7<br>(2.4%) | 0.02<br>(0.03%) |
| **96** | 38.5<br>(43.2%) | 16.5<br>(21.7%) | 7.7<br>(10.1%) | 11.0<br>(14.5%) | 4.7<br>(6.2%) | 1.5<br>(1.9%) | 0.6<br>(0.8%) | 1.2<br>(1.6%) | 0.03<br>(0.04%) |
| **250** | 70.2<br>(44.5%) | 57.8<br>(42.8%) | 7.7<br>(5.7%) | 3.8<br>(2.8%) | 3.6<br>(2.6%) | 0.6<br>(0.5%) | 1.2<br>(0.9%) | N/A | N/A |
| **70** | 52.3<br>(48.1%) | 19.8<br>(21.1%) | 13.3<br>(14.2%) | 6.4<br>(6.8%) | 3.4<br>(3.6%) | 3.8<br>(4.0%) | 0.7<br>(0.8%) | 1.1<br>(1.2%) | 0.1<br>(0.1%) |
| **Overall** | **36.5**<br>**(40.1%)** | **22.2**<br>**(24.2%)** | **7.5**<br>**(10.1%)** | **11.0**<br>**(15.3%)** | **3.3**<br>**(4.7%)** | **2.1**<br>**(3.0%)** | **0.9**<br>**(1.0%)** | **1.1**<br>**(1.7%)** | **0.1**<br>**(0.1%)** |

**Supplemental Table 3.**
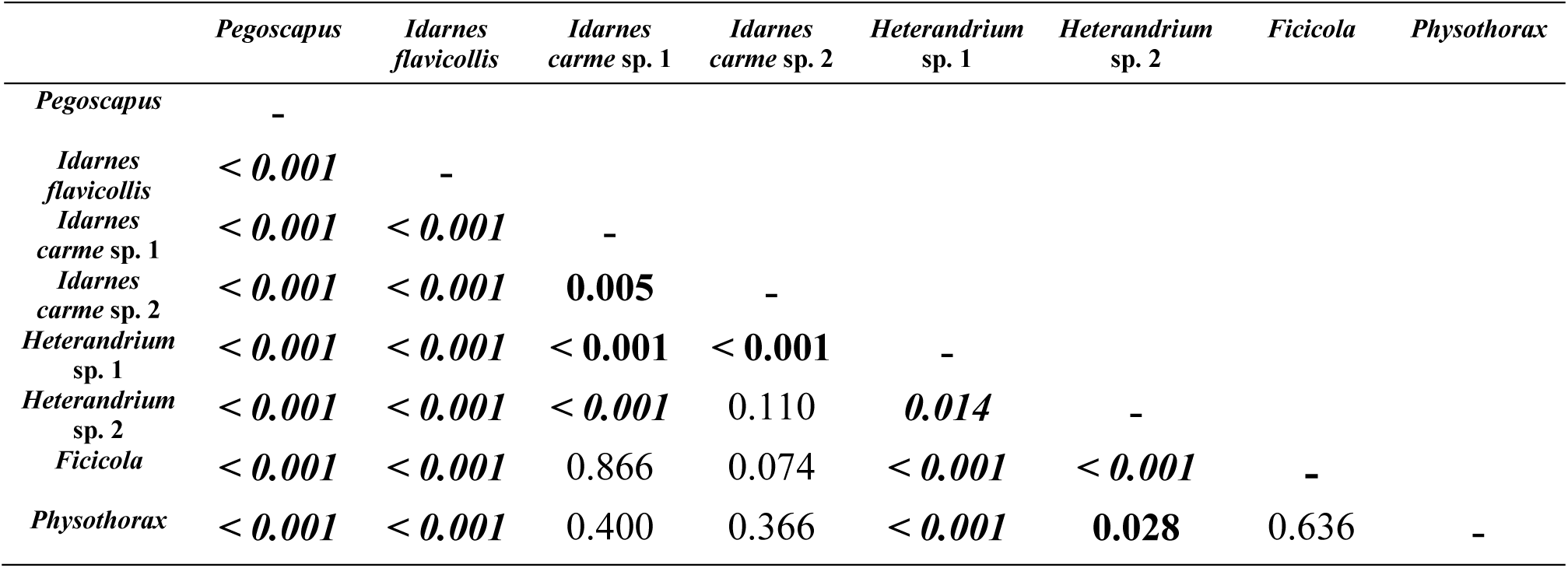
Matrix result of exact tests (using poisson.test in R) comparing numbers of juvenile *Parasitodiplogaster* nematodes per host between pairs of pollinator and NPFW species associated with *F. petiolaris. P*-values are in bold where the row species has a significantly higher nematode load, and in bold italics where the column species has the higher load.

**Supplemental Table 4.** Nematode infection of successfully dispersed NPFWs sampled from receptive *F. petiolaris* figs.

|  | **Wasps**<br>**Dissected** | **Percent**<br>**Infected** | **Average**<br>**Nematodes**<br>**per Host** | **Median**<br>**Nematodes**<br>**per Host** | **Maximum**<br>**Nematodes**<br>**per Host** |
| --- | --- | --- | --- | --- | --- |
| *Idarnes flavicollis* | 111 | 3.6% | 1.25 | 1 | 2 |
| *Idarnes carme* sp. 1 | 86 | 1.2% | 1 | 1 | 1 |
| *Idarnes carme* sp. 2 | 71 | 2.8% | 1 | 1 | 1 |
| *Heterandrium* sp. 1 | 3 | N/A | N/A | N/A | N/A |
| *Heterandrium* sp. 2 | 3 | N/A | N/A | N/A | N/A |
| *Physothorax* | 1 | N/A | N/A | N/A | N/A |

**Supplemental Table 5.**
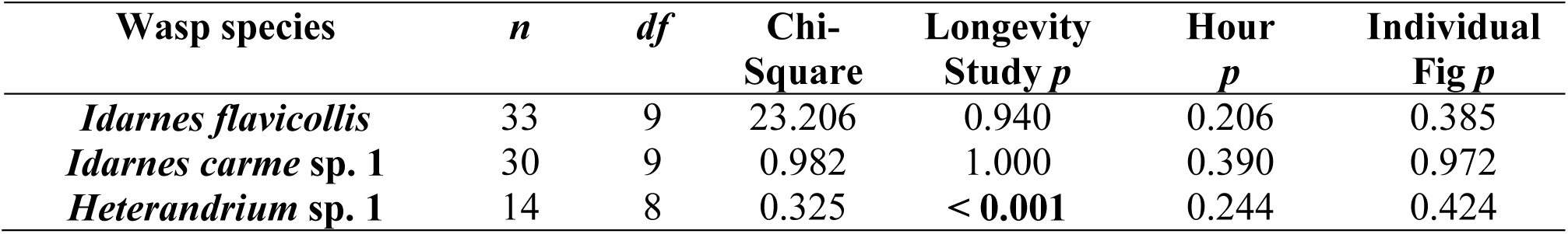
Details of the Generalized Linear Model (GLM) analyses conducted for a NPFW controlled longevity trials in *F. petiolaris* from 50 controlled figs total (29 infested, 21 uninfested). These tables represent individual NPFW species that experienced adequate infection rates to evaluate meaningful longevity analyses (> 10 individuals infected). We did not observe significant NPFW longevity reductions due to number of nematodes involved in an infective event in these studies. Likewise, we were unable to find a significant difference between the survivorship curves of infected and uninfected individuals in any of these three species (Wilcoxon test, *p*-values = 0.180, 0.838, and 0.299 respectively). In these models the response variable *number of nematodes extracted per wasp host* was analyzed against the predictor variables *longevity study, hour*, and *individual fig*. Displayed here are the model details (*n, df*, test Chi-Square value, and predictor variable *p*-values) for each of the three abundant NPFW species studied in this section. Significant *p-values* (*p* < 0.05) are displayed in bold.

| Wasp species | *n* | *df* | Chi-Square | Longevity Study *p* | Hour *p* | Individual Fig *p* |
| --- | --- | --- | --- | --- | --- | --- |
| *Idarnes flavicollis* | 33 | 9 | 23.206 | 0.940 | 0.206 | 0.385 |
| *Idarnes carme* sp. 1 | 30 | 9 | 0.982 | 1.000 | 0.390 | 0.972 |
| *Heterandrium* sp. 1 | 14 | 8 | 0.325 | < **0.001** | 0.244 | 0.424 |

**Supplemental Table 6.** Details of the ANOVA analyses conducted for the *F. petiolaris* controlled longevity trial for NPFWs. We did not observe a significant effect of nematode infection on the hour in which infected NPFW died in these trials. In these models the response variable *hour* was analyzed against the predictor variables *longevity study, individual fig*, and *number of nematodes extracted per wasp host*. Displayed here are the model details (*n, df, F, r*^*2*^, and predictor variable *p*-values) for each of the three non-pollinating wasp species studied in this section. Significant *p-values* (*p* < 0.05) are displayed in bold.

| Wasp species | *n* | *df* | *F* | *r*<sup>2</sup> | Longevity<br>Study<br>*p* | Individual<br>Fig<br>*p* | Number of<br>Nematodes<br>per Wasp<br>Host *p* |
| --- | --- | --- | --- | --- | --- | --- | --- |
| *Idarnes flavicollis* | 33 | 9 | 5.411 | 0.679 | 0.167 | 0.189 | 0.415 |
| *Idarnes carme* sp. 1 | 30 | 9 | 4.342 | 0.661 | 1.000 | **0.002** | 0.107 |
| *Heterandrium* sp. 1 | 14 | 8 | 1.340 | 0.682 | 0.171 | 0.419 | 0.315 |

**Supplemental Table 7.** Nematode infection of pollinator (*Pegoscapus*) and NPFWs (three distinct *Idarnes* species) emerging from mature, nematode-infested figs *F. popenoei* in Panama.

| Wasp Species | Wasps Dissected | Percent Infected | Average Nematodes per Host | Median Nematodes per Host | Maximum Nematodes per Host |
| --- | --- | --- | --- | --- | --- |
| *Pegoscapus* | 78 | 37% | 3.17 | 2 | 11 |
| *Idarnes flavicollis* | 143 | 41% | 3.49 | 2 | 17 |
| *Idarnes carme* | 118 | 6% | 1.43 | 1 | 4 |
| *Idarnes inserta* | 9 | 0% | N/A | N/A | N/A |

**Supplemental Table 8.** Details of the Generalized Linear Model (GLM) analyses conducted for the *F. popenoei* controlled longevity trial carried out on Barro Colorado Island, Panama in July 2017. *F. popenoei* was chosen here for a longevity trial due to similar wasp population dynamics when compared to *F. petiolaris* (mostly single pollinating foundresses and many NPFW). Here, our data suggests that nematode infection significantly limits the longevity of one NPFW, but not the pollinator. In these models the response variable *number of nematodes extracted per wasp host* was analyzed against the predictor variables *individual fig* and *hour*. Displayed here are the model details (*n, df*, test Chi-Square value, and predictor variable *p*-values) for each of the three species studied in this section. Significant *p-values* are displayed in bold. However, through these efforts, we were unable to find a significant difference in the survivorship curves of infected and uninfected *Pegoscapus* or *Idarnes flavicollis* wasps (Wilcoxon test, *p* = 0.889 and 0.219 respectively).

| Wasp Species | *n* | *df* | Chi-Square | Individual<br>Fig *p* | Hour<br>*p* |
| --- | --- | --- | --- | --- | --- |
| *Pegoscapus* | 29 | 6 | 10.013 | 0.187 | 0.670 |
| *Idarnes flavicollis* | 59 | 8 | 42.105 | **0.002** | **< 0.001** |
| *Idarnes carme* | 7 | 3 | 3.670 | 0.561 | 0.085 |

**Supplemental Table 9.** Details of the ANOVA analyses conducted for the *F. popenoei* controlled longevity trial in Panama. Here, our data suggests that the hour in which two of the NPFWs died in this study were significantly influenced by the number of nematodes infecting them, but not for the pollinator wasp. In these models the response variable *hour* was analyzed against the predictor variables *individual fig*, and *number of nematodes extracted per wasp host*. Displayed here are the model details (*n, df, F, r*^*2*^, and predictor variable *p*-values) for each of the three non-pollinating wasp species studied in this section. Significant *p-values* (*p* < 0.05) are displayed in bold.

| Wasp Species | *n* | *df* | *F* | *r*<sup>2</sup> | Individual<br>Fig *p* | Number of<br>Nematodes<br>per Wasp<br>Host *p* |
| --- | --- | --- | --- | --- | --- | --- |
| *Pegoscapus* | 29 | 6 | 2.782 | 0.431 | **0.030** | 0.820 |
| *Idarnes flavicollis* | 59 | 8 | 3.165 | 0.336 | **0.017** | **0.016** |
| *Idarnes carme* | 7 | 3 | 16.600 | 0.943 | **0.031** | **0.032** |

## Supplemental Figures

**Supplemental Figure 1.**
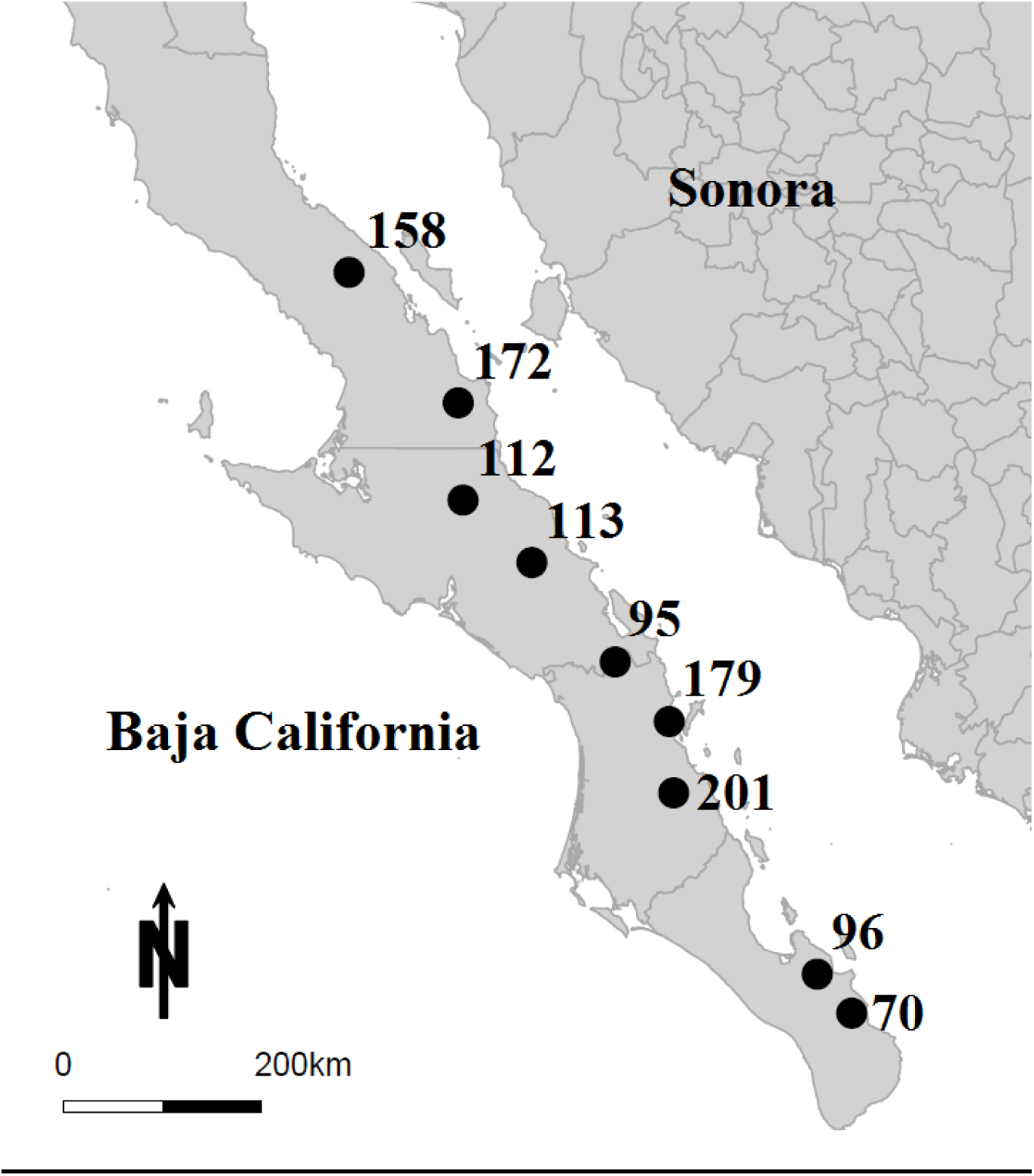
Map of *Ficus petiolaris* study sites in Baja California and Baja California Sur Mexico.

## Notes

### Competing Interest Statement

The authors have declared no competing interest.

